# Prevalence of beta-lactam drug-resistance genes in commensal *Escherichia coli* contaminating ready-to-eat lettuce

**DOI:** 10.1101/824516

**Authors:** Ningbo Liao, Julia Rubin, Yuan Hu, Hector A. Ramirez, Clarissa Araújo Borges, Biao Zhou, Yanjun Zhang, Ronghua Zhang, Jianmin Jiang, Lee W. Riley

**Affiliations:** School of Public Health, Division of Infectious Diseases and Vaccinology, University of California, Berkeley, California 94720, USA; Department of Nutrition and Food Safety, Zhejiang Provincial Center for Disease Control and Prevention, Hangzhou 310006, China

## Abstract

The objective of this study was to evaluate the prevalence of antibiotic resistance and beta-lactam drug resistance genes in *Escherichia coli* isolated from ready-to-eat lettuce, obtained from local supermarkets in Northern California. Bags of lettuce were purchased from 4 chain supermarkets during three different periods—Oct 2018–Jan 2019, Feb 2019–Apr 2019 and May 2019–July 2019. From 91 packages of lettuce, we recovered 34 *E. coli* isolates from 22 (24%) lettuce samples. All *E. coli* isolates were genotyped by multilocus sequence typing (MLST), and we found 15 distinct sequence types (STs). Five of these genotypes (ST2819, ST4600, ST2432, ST1198 and ST5143) have been reported to cause infection in humans. Twenty (59%) *E. coli* isolates were found resistant to at least one of the antibacterial drugs. They included resistance to ampicillin (AMP, 85%) and ampicillin/sulbactam (SAM, 50%), cefoxitin (FOX, 40%) and cefuroxime (CXM, 35%). We found 8 (40%) of 20 beta-lactam resistant *E. coli* isolates to carry *bla*_CTX-M_; 5 (25%) tested positive for *bla*_SHV_, while only 4 (20%) tested positive for *bla*_TEM_. Additionally, we identified a class A broad-spectrum beta-lactamase SED-1 gene, *bla*_SED_, reported by others in *Citrobacter sedlakii* isolated from bile of a patient. This study found that a large proportion of fresh lettuce carry beta-lactam drug-resistant *E. coli*, which could serve as a reservoir for drug resistance genes that could potentially enter pathogens to cause human infections.

## INTRODUCTION

Foodborne illnesses caused by various microorganisms including bacteria are major concern to public health in both developed and developing countries (1). Antimicrobial drugs have played an indispensable role in the treatment of foodborne infectious diseases, but the frequent clinical use of antimicrobial agents has resulted in the selection of antimicrobial drug-resistant (AMR) bacterial strains. However, factors that contribute to the widespread dissemination of AMR bacterial strains in community are not limited to just the human clinical use of antimicrobial agents (2).

Foods may contribute as an important source of antibiotic resistance genes found in the human intestine. In particular, fresh produce carry saprophytic bacteria that harbor drug-resistance genes, which can enter the human gut when it is eaten uncooked (3, 4). Green leafy vegetables are frequently colonized with Gram-negative bacteria, especially by members of *Pseudomonadaceae* and *Enterobacteriaceae* (5). Several foodborne disease outbreaks caused by lettuce contaminated with recognized pathogens (*E. coli* O157:H7, Salmonella) have been increasingly reported in the United States in the last 10 years (6, 7, 8, 9, 10).

The primary reservoirs of these enteric pathogens are the food animal intestines, and the green leafy vegetables are believed to become contaminated by food animal manure. Thus, commensal *E. coli* in manure from such animals could also contaminate green leafy vegetables. The objective of this study was, therefore, to determine the prevalence of *E. coli* contamination of packaged, ready-to-eat lettuce obtained from chain supermarkets in a Northern California community, and assess the frequency of their antimicrobial drug resistance and beta-lactam drug resistance gene carriage.

## MATERIALS AND METHODS

In this study, ready-to-eat lettuce packages were purchased from chain supermarkets located within a 7-mile vicinity of a university community in Northern California during three different periods—Oct 2018–Jan 2019, Feb 2019–Apr 2019 and May 2019–July 2019. For each lettuce package, 25 g of lettuce were placed in a UV-pretreated polyethylene bag containing 50 ml of sterile phosphate-buffered saline (PBS; pH 7.4). The lettuce was incubated in PBS for 30 to 60 min at room temperature after brief kneading. The PBS wash was then transferred to a 50-ml conical tube and centrifuged at 12,000 g for 5 min at room temperature. The resulting pellet was resuspended in 2 ml of PBS, and 1 ml of it was saved in a 10% glycerol stock. The rest of the suspension (10μL) was cultured on a MacConkey agar plate to isolate Gram-negative bacteria and presumptively identified by lactose fermentation and indole test as *E. coli*.

Five colonies recovered on plate from each sample were randomly picked and subtyped by the enterobacterial repetitive intergenic consensus (ERIC)-PCR assay according to standard procedures (11). Single colonies from tryptic soy agar plates were selected and inoculated into 2 ml tryptic soy broth and incubated in a shaking incubator for 15 h at 37°C. The 1-mL aliquots of grown cultures were centrifuged, and the pellets were collected to isolate DNA as described by Yamaji et al. (12). The five colonies that had identical ERIC electrophoretic banding patterns by visual inspection were considered to belong to the same clonal group, and one of these colonies was selected for further analysis by multilocus sequence typing (MLST). If samples contained more than one distinct electrophoretic banding pattern by ERIC-PCR typing, all strains were included in the MLST analysis (12, 13). The allelic number and the corresponding genotype number were designated by the curator of the MLST based on the seven-gene scheme described at website http://mlst.ucc.ie/mlst/dbs/Ecoli (13, 14).

All isolates from lettuce samples were assessed for susceptibility to ampicillin (AMP), cefoxitin (FOX), cefotaxime (CTX), cefuroxime (CXM), nitrofurantoin (NIT), ceftriaxone (CRO), ampicillin/sulbactam (SAM), ceftazidime (CAZ), fomycin glucose (FOS), gentamicin (CN), trimethoprim-sulfamethozaxole (TMP-SMX), and nalidixic acid (NA) by the standard disc diffusion assay (15).

The analysis of beta-lactamase genes was performed according to previously published reports (16, 17). Primers (Table S1) used for PCR amplification of the beta-lactams genes (*bla*_TEM_, *bla*_SHV_, *bla*_CXT-M_ and *bla*_OXA_) were acquired from Sigma Aldrich, USA. The PCR amplification was carried out with 5 μL of DNA template (100-200 ng/μL) in 25 μL PCR mixture. Each reaction mixture contained 1 μL of each, reverse and forward primer (10 μM), PCR grade water 1.5 μL, and Taq PCR master mix (Bio-Rad, USA) 16.5 μL. Amplification reactions were accomplished with a DNA thermo-cycler (Bio-Rad) as follows: initial denaturation for 30 seconds at 95°C, followed by 30 cycles of denaturation at 95°C for 30 seconds, annealing for 60 seconds (59°C, 60°C, 61°C and 65°C for *bla*_TEM_, *bla*_SHV_, *bla*_CXT-M_ and *bla*_OXA_, respectively). After final extension for 5 min at 72°C, amplified samples were subjected to gel electrophoresis with QIAxcel advance system (QIAGEN, USA) (16). DNA sequences of 400 bp or more for *bla*_TEM_, *bla*_SHV_ and *bla*_CXT-M_ were visually inspected, aligned, and compared against sequences in GenBank with BLAST (https://blast.ncbi.nlm.nih.gov/Blast.cgi). Statistical analysis was performed by Fisher’s exact test with significance set at *p* ≤0.05.

## RESULTS AND DISCUSSION

Ninety-one lettuce packages were purchased from chain supermarkets located within a 7-mile vicinity of a university community in Northern California. They included organic (n=47) and non-organic (n=44) lettuce samples (Table S2). From 91 lettuce samples, we recovered 34 *E. coli* isolates from 22 (24%) samples. *E. coli* was isolated from 18 (38%) organic and 9 (20%) non-organic lettuce packages (*p*=0.051, Fisher’s exact test, 1-tailed). By MLST analysis, among 34 *E. coli* isolates, 24 were assigned to 15 unique STs (Table 1). Two genotypes (ST8951 and ST5154) were commonly detected in the lettuce samples. These genotypes have been reported in deciduous forest soil and cattle, but not from human infections in the MLST database (http://enterobase.warwick.ac.uk). We found strains ST2819, ST4600, ST2432, ST5143 and ST1198, which have been reported in other regions of the world including Japan (Fukuoka), USA (Nashville), China (Hangzhou and Shanghai), and Korea. They have been isolated from cases of human infection, as noted in the MLST database. As shown in Table 1, *E. coli* strains ST4600 and ST1198 have been reported as causes of diarrhea and urinary tract infection in USA and Korea, respectively.

**Table 1.** *E. coli* isolates belonging to the 15 MLST genotypes found in lettuce samples

| ST(total no. of isolates typed ) | ST Complex | In MLST datas (http://enterobase.warwick.ac.uk/species/ecoli/search_strains?query=st_search) |  |  |  |
| --- | --- | --- | --- | --- | --- |
|  |  | Source | Collection year | Location (Country or City) | Path/Nonpath |
| ST8951(4) | None | Deciduous Forest; Deciduous Forest Soil; Soil | 2015 | None | None |
| ST2819(2) | None | human | 2015 | Japan(Fukuoka) | Pathogen (Diarrhoea) |
| ST5154(3) | None | Cattle | 2017 | Africa(Angola) | None |
| ST6374(2) | None | None | None | None | None |
| ST9285(1) | None | Poultry | 2016 | Africa(Gambia) | None |
| ST5143(2) | None | Faeces | 2013 | China(Shanghai) | Pathogen |
| ST2432(2) | None | human | 2011 | China(Hangzhou) | Pathogen |
| ST5351(1) | ST568Cplx | None | None | None | None |
| ST3567(1) | None | human | 2012 | Netherlands | None |
| ST2625(1) | ST14Cplx | human | 2015 | USA(Seattle) | Pathogen |
| ST4277(1) | None | Oryzomys sp | 2015 | Mexico | None |
| ST4600(1) | None | Human | 2011 | USA(Nashville) | Pathogen (Diarrhoea) |
| ST3222(1) | ST648Cplx | human | 2008 | France(Besancon) | None |
| ST1198(1) | None | human | 2007 | Korea | Pathogen (UTI) |
| ST6376(1) | None | None | None | None | None |
| unknown(10) |  | None | None | None | None |

Antibiotic susceptibility testing showed that most of the *E. coli* isolates were resistant to at least one of the tested antibiotics (Table 2). Among 34 *E. coli* isolates 20 (59%) were resistant to one or more antibiotics; 17 (85%) were resistant to ampicillin (AMP), 10 (50%) to ampicillin/sulbactam (SAM), 8 (40%) to cefoxitin (FOX), 7 (35%) to cefuroxime (CXM), and only 1 (5%) to nalidixic acid (NA). Nine (45%) of these isolates were multidrug resistant (MDR), defined as resistance to three or more classes of drugs (Table S3). Beta-lactam-resistant strains were examined for beta-lactamase genes by PCR. Eight (40%) of 20 beta-lactam resistant *E. coli* strains contained *bla*_CTX-M_, 5 (25%) tested positive for *bla*_SHV_, and 4 (20%) tested positive for *bla*_TEM_. The high frequency of *E. coli* strains carrying *bla*_CTX-M_ is surprising, since most beta-lactam resistant clinical isolates of *E. coli* carry *bla*_TEM_ and/or *bla*_SHV_ (17, 18, 19). Of 9 MDR *E. coli* isolates that showed resistance to ampicillin, ampicillin/sulbactam, cefoxitin and cefuroxime, 8 tested positive for *bla*_TEM_, *bla*_CTX-M_ or *bla*_SHV_ genes. CTX-M beta-lactamases are the most common extended-spectrum beta-lactamases (ESBL) expressed by clinical isolates of *E. coli*, which are increasingly reported in extraintestinal infections from many regions of the world, especially in healthcare settings (20, 21, 22).

**Table 2.**
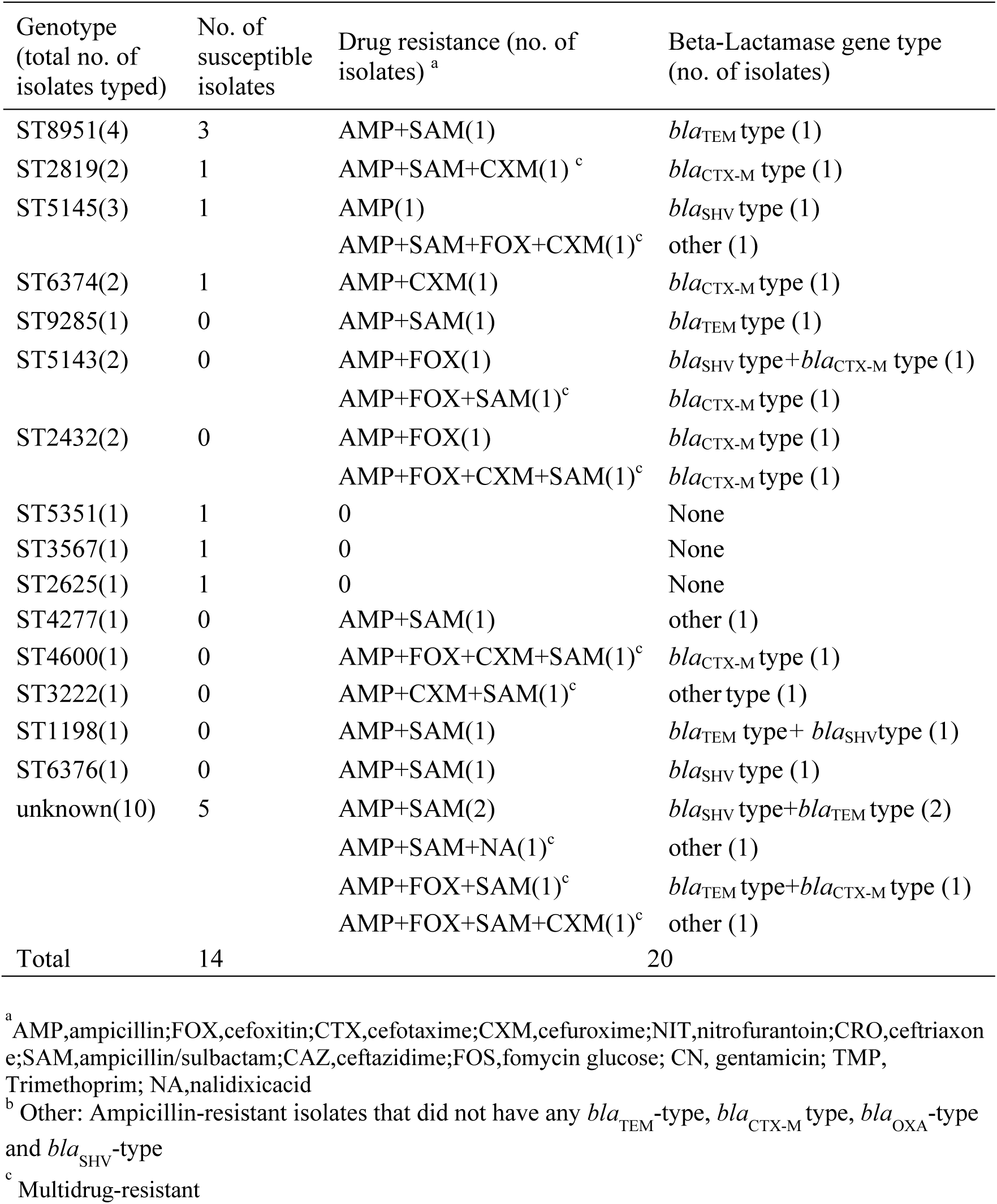
Beta-Lactamase gene types identified among antibiotic-resistant *E. coli* isolates from lettuces.

| Genotype<br>(total no. of<br>isolates typed) | No. of<br>susceptible<br>isolates | Drug resistance (no. of<br>isolates)<sup>a</sup> | Beta-Lactamase gene type<br>(no. of isolates) |
| --- | --- | --- | --- |
| ST8951(4) | 3 | AMP+SAM(1) | *bla*<sub>TEM</sub> type (1) |
| ST2819(2) | 1 | AMP+SAM+CXM(1)<sup>c</sup> | *bla*<sub>CTX-M</sub> type (1) |
| ST5145(3) | 1 | AMP(1) | *bla*<sub>SHV</sub> type (1) |
|  |  | AMP+SAM+FOX+CXM(1)<sup>c</sup> | other (1) |
| ST6374(2) | 1 | AMP+CXM(1) | *bla*<sub>CTX-M</sub> type (1) |
| ST9285(1) | 0 | AMP+SAM(1) | *bla*<sub>TEM</sub> type (1) |
| ST5143(2) | 0 | AMP+FOX(1) | *bla*<sub>SHV</sub> type+*bla*<sub>CTX-M</sub> type (1) |
|  |  | AMP+FOX+SAM(1)<sup>c</sup> | *bla*<sub>CTX-M</sub> type (1) |
| ST2432(2) | 0 | AMP+FOX(1) | *bla*<sub>CTX-M</sub> type (1) |
|  |  | AMP+FOX+CXM+SAM(1)<sup>c</sup> | *bla*<sub>CTX-M</sub> type (1) |
| ST5351(1) | 1 | 0 | None |
| ST3567(1) | 1 | 0 | None |
| ST2625(1) | 1 | 0 | None |
| ST4277(1) | 0 | AMP+SAM(1) | other (1) |
| ST4600(1) | 0 | AMP+FOX+CXM+SAM(1)<sup>c</sup> | *bla*<sub>CTX-M</sub> type (1) |
| ST3222(1) | 0 | AMP+CXM+SAM(1)<sup>c</sup> | other type (1) |
| ST1198(1) | 0 | AMP+SAM(1) | *bla*<sub>TEM</sub> type+ *bla*<sub>SHV</sub>type (1) |
| ST6376(1) | 0 | AMP+SAM(1) | *bla*<sub>SHV</sub> type (1) |
| unknown(10) | 5 | AMP+SAM(2) | *bla*<sub>SHV</sub> type+*bla*<sub>TEM</sub> type (2) |
|  |  | AMP+SAM+NA(1)<sup>c</sup> | other (1) |
|  |  | AMP+FOX+SAM(1)<sup>c</sup> | *bla*<sub>TEM</sub> type+*bla*<sub>CTX-M</sub> type (1) |
|  |  | AMP+FOX+SAM+CXM(1)<sup>c</sup> | other (1) |
| Total | 14 |  | 20 |

<sup>a</sup> AMP, ampicillin; FOX, ceftiofur; CTX, cefotaxime; CXM, cefuroxime; NIT, nitrofurantoin; CRO, ceftriaxone; SAM, ampicillin/sulbactam; CAZ, ceftazidime; FOS, fosfomicin; CN, gentamicin; TMP, Trimethoprim; NA, nalidixic acid
<sup>b</sup> Other: Ampicillin-resistant isolates that did not have any *bla*<sub>TEM</sub>-type, *bla*<sub>CTX-M</sub> type, *bla*<sub>OXA</sub>-type and *bla*<sub>SHV</sub>-type
<sup>c</sup> Multidrug-resistant

To our surprise, in one strain (ST3222), we found a class A broad-spectrum beta-lactamase SED-1 gene (Table 3), *bla*_SED_, reported by others to have been found in *Citrobacter sedlakii* in the bile fluid of a patient (23, 24). This observation suggests that some of these beta-lactamase genes are horizontally transferred among saprophytic Gram-negative bacteria found in the environment, and that commensal *E. coli* from mammalian intestine contaminating lettuce could acquire them. Such genes can then enter the human intestine via ingestion of such contaminated food product eaten uncooked.

**Table 3.** Genetic analysis of the beta-Lactamase -producing *E.coli* in lettuces

| Sequence type | Strain | Lettuce type | Size (bp) of sequence DNA fragment | Position (bp) of sequence on matched sequence | Matched sequences in NCBI database (organism of origin,accession no,sequence length [bp], % of Matches) |
| --- | --- | --- | --- | --- | --- |
| ST2819 | E04270401 | Organic | 446 | 33-478 | *bla*<sub>CTX-M</sub> gene, *Citrobacter sp.*, KJ918511.1, 525, 99 |
| ST4600 | E06060204 | Organic | 425 | 53-478 | *bla*<sub>CTX-M</sub> gene, *Citrobacter sp.*, KJ918511.1, 525, 99 |
| ST2432 | E06200301 | Organic | 433 | 45-478 | *bla*<sub>CTX-M</sub> gene, *Citrobacter sp.*, KJ918511.1, 525, 99 |
| ST3222 | E06200202 | Organic | 455 | 270-725 | *bla*<sub>SED</sub> gene, *Citrobacter sedlakii*, NG_049979.1,888, 98.68 |
| ST1198 | E50802 | Organic | 663 | 122-784 | *bla*<sub>SHV-11</sub> gene, *Escherichia coli*, MF402896.1, 861, 99.55 |
| ST5143 | E06270403 | Non-organic | 653 | 131-784 | *bla*<sub>SHV-11</sub> gene, *Escherichia coli*, MF402896.1, 861, 99.55 |
| unknown | E05180501 | Non-organic | 388 | 261-648 | *bla*<sub>TEM</sub> gene, *Acinetobacter baumannii*, MK764360.1, 732, 100 |
| unknown | E06270303 | Organic | 433 | 212-768 | *bla*<sub>SHV-11</sub> gene, *Citrobacter freundii*, MF402898.1, 861, 89.59 |

The limitation of this study is that we analyzed the resistant *E. coli* strains for only beta-lactamase genes. Since many of the strains were resistant to other classes of drugs, they are likely to carry other drug-resistance genes. Hence, our study underestimates the magnitude of these lettuce samples contributing to the spread of these genes into the human intestine.

## CONCLUSION

A large proportion of retail ready-to-eat lettuce samples contained *E. coli*, many of which were resistant to commonly-used beta-lactam antibiotics. Although the predominant STs (ST8951 and ST5154) were not pathogens, other genotypes ST2819, ST4600, ST2432, ST1198 and ST5143, which were multidrug-resistant, have been isolated from human infections in several regions of the world. A surprisingly large proportion of the beta-lactam-resistant strains carried an ESBL beta-lactamase gene *bla*_CTX-M_. While green-leafy vegetables could serve as sources of recognized enteric pathogens, they may also serve as a common vector for drug-resistance genes that ultimately enter human pathogens.

## ACKNOWLEDGMENT

This work was funded by the U.S. Centers for Disease Control and Prevention program to combat antibiotic resistance under BAA number 200-2016-91939, and the National Natural Science Foundation of China (#31301715), the Program of Health and Medicine Science in Zhejiang Province, China (#2017KY292), and the National Health Commission Foundation of the People’s Republic of China (WKJ-ZJ-1917).

